## Supplementary Table 1 for "Insights into the RNA virome of the corn leafhopper *Dalbulus maidis*, a major emergent threat of Maize in Latin America"

**Supplementary Table 1.** NCBI publicly available datasets used for virus discovery. Total virus reads and per million (RPM) are based on mapping results using Bowtie.2 with standard parameters against the genome consensus sequences of Chicharrita del maiz orthomyxo-like virus 1 (ChMOMLV), Chicharrita del maiz beny-like virus 1 (ChMBLV), Chicharrita del maiz rhabdo-like virus 1 (ChMRV), Chicharrita del maiz iflavirus 1 (ChMIfV1), Chicharrita del maiz iflavirus 2 (ChMIfV2), and Chicharrita del maiz bunyan-like virus 1 (ChMBV).

| **Run** | **BioProject** | **BioSample** | **Experiment** | **Country** | **Total library reads** | **ChMOMLV reads** | **ChMOMLV RPM** | **ChMBLV reads** | **ChMBLV RPM** | **ChMRV reads** | **ChMRV RPM** | **ChMIfV1**  **reads** | **ChMIfV1 RPM** | **ChMIfV2 reads** | **ChMIfV2 RPM** | **ChMBV reads** | **ChMBV RPM** |
| --- | --- | --- | --- | --- | --- | --- | --- | --- | --- | --- | --- | --- | --- | --- | --- | --- | --- |
| SRR8248833 | PRJNA505618 | SAMN10430478 | SRX5066649 | Argentina | 44568016 | **2** | 0,0 | 0 | 0,0 | **6720** | **150,8** | **151** | **3,4** | 0 | 0,0 | 0 | 0,0 |
| SRR10334087 | PRJNA562189 | SAMN13091747 | SRX7044228 | USA | 210894690 | 0 | 0,0 | 0 | 0,0 | 0 | 0,0 | 0 | 0,0 | **10846** | **51,4** | 0 | 0,0 |
| SRR15166998 | PRJNA562189 | SAMN20251994 | SRX11474353 | USA | 45974618 | **3020** | **65,7** | **13192** | **286,9** | 0 | 0,0 | **16114** | **350,5** | **680** | **14,8** | 0 | 0,0 |
| SRR1821981 | PRJNA272239 | SAMN03341953 | SRX893575 | USA | 697217 | **172** | **246,7** | 0 | 0,0 | 0 | 0,0 | 0 | 0,0 | 0 | 0,0 | **540** | **774,5** |
| SRR5010611 | PRJNA318849 | SAMN06009808 | SRX2341563 | Brazil | 44939338 | **949** | **21,1** | 0 | 0,0 | 0 | 0,0 | **19631** | **436,8** | **1114** | **24,8** | **363** | **8,1** |
| SRR5010612 | PRJNA318849 | SAMN06009807 | SRX2341564 | Brazil | 45008074 | **397** | **8,8** | 0 | 0,0 | 0 | 0,0 | **11422** | **253,8** | **180** | **4,0** | **388** | **8,6** |
| SRR5010613 | PRJNA318849 | SAMN06009806 | SRX2341565 | Brazil | 43120958 | 0 | 0,0 | 0 | 0,0 | 0 | 0,0 | **9596** | **222,5** | **619** | **14,4** | **573** | **13,3** |
| SRR11822333 | PRJNA631706 | SAMN14890635 | SRX8372961 | USA | 36054352 | 0 | 0,0 | 0 | 0,0 | 0 | 0,0 | 0 | 0,0 | **10732** | **297,7** | 0 | 0,0 |
| SRR11822334 | PRJNA631706 | SAMN14890635 | SRX8372960 | USA | 42673870 | 0 | 0,0 | 0 | 0,0 | 0 | 0,0 | 0 | 0,0 | **23298** | **546,0** | 0 | 0,0 |
| SRR11822335 | PRJNA631706 | SAMN14890505 | SRX8372959 | USA | 41853828 | 0 | 0,0 | 0 | 0,0 | 0 | 0,0 | 0 | 0,0 | **6677** | **159,5** | 0 | 0,0 |
| SRR11822336 | PRJNA631706 | SAMN14890505 | SRX8372958 | USA | 51372620 | 0 | 0,0 | 0 | 0,0 | 0 | 0,0 | 0 | 0,0 | **4168** | **81,1** | 0 | 0,0 |
| SRR11822337 | PRJNA631706 | SAMN14890505 | SRX8372957 | USA | 40536434 | 0 | 0,0 | 0 | 0,0 | 0 | 0,0 | 0 | 0,0 | **10398** | **256,5** | 0 | 0,0 |
| SRR11822338 | PRJNA631706 | SAMN14890505 | SRX8372956 | USA | 24793552 | 0 | 0,0 | 0 | 0,0 | 0 | 0,0 | 0 | 0,0 | **12651** | **510,3** | 0 | 0,0 |
| SRR11822339 | PRJNA631706 | SAMN14890446 | SRX8372955 | USA | 34075044 | 0 | 0,0 | 0 | 0,0 | 0 | 0,0 | 0 | 0,0 | **10623** | **311,8** | 0 | 0,0 |
| SRR11822340 | PRJNA631706 | SAMN14890446 | SRX8372954 | USA | 45038780 | 0 | 0,0 | 0 | 0,0 | 0 | 0,0 | 0 | 0,0 | **4127** | **91,6** | 0 | 0,0 |
| SRR11822341 | PRJNA631706 | SAMN14890756 | SRX8372953 | USA | 46079864 | 0 | 0,0 | 0 | 0,0 | 0 | 0,0 | 0 | 0,0 | **24639** | **534,7** | 0 | 0,0 |
| SRR11822342 | PRJNA631706 | SAMN14890756 | SRX8372952 | USA | 36986548 | 0 | 0,0 | 0 | 0,0 | 0 | 0,0 | 0 | 0,0 | **14839** | **401,2** | 0 | 0,0 |
| SRR11822343 | PRJNA631706 | SAMN14890756 | SRX8372951 | USA | 42869668 | 0 | 0,0 | 0 | 0,0 | 0 | 0,0 | 0 | 0,0 | **37763** | **880,9** | 0 | 0,0 |
| SRR11822344 | PRJNA631706 | SAMN14890756 | SRX8372950 | USA | 38197304 | 0 | 0,0 | 0 | 0,0 | 0 | 0,0 | 0 | 0,0 | **23300** | **610,0** | 0 | 0,0 |
| SRR11822345 | PRJNA631706 | SAMN14890635 | SRX8372949 | USA | 48481712 | 0 | 0,0 | 0 | 0,0 | 0 | 0,0 | 0 | 0,0 | **16570** | **341,8** | 0 | 0,0 |
| SRR11822346 | PRJNA631706 | SAMN14890635 | SRX8372948 | USA | 42292438 | 0 | 0,0 | 0 | 0,0 | 0 | 0,0 | 0 | 0,0 | **11644** | **275,3** | 0 | 0,0 |
| SRR11822347 | PRJNA631706 | SAMN14890446 | SRX8372947 | USA | 43790788 | 0 | 0,0 | 0 | 0,0 | 0 | 0,0 | 0 | 0,0 | **2662** | **60,8** | 0 | 0,0 |
| SRR11822348 | PRJNA631706 | SAMN14890446 | SRX8372946 | USA | 22991808 | 0 | 0,0 | 0 | 0,0 | 0 | 0,0 | 0 | 0,0 | **10249** | **445,8** | 0 | 0,0 |
