## Supplementary material for "Insights into the RNA virome of the corn leafhopper *Dalbulus maidis*, a major emergent threat of Maize in Latin America": Table 1

**Table 1. Summary of the *Dalbulus maidis* viruses identified and genomic features.**

| **Virus name/abbreviation** | **Accession number** | **Genome/ segment lenght (nt)** | **Protein ID/lenght (aa)** | **Highest scoring virus- protein / *E*-value / query coverage% / identity % (Blast P)** |
| --- | --- | --- | --- | --- |
| Chicharrita del maíz beny-like virus/ ChMBLV | BK068250 | 6,887 | Polyprotein / 2,003  structural P / 224 | DuV2-polyprotein / 0.0 / 70 / 46.61  DuV2-structural P / 8e-07 / 53 / 29.75 |
| Chicharrita del maíz bunyan-like virus/ChMBV | BK068254  BK068255 | Segment L 6,651  Segment S 1,281 | RdRp / 2,166  NP / 327 | HbIV1-RdRp / 0.0 / 99 / 49.84  HbIV1-NP / 8e-53 / 81 / 38.01 |
| Chicharrita del maíz iflavirus 1/ChMIfV1 | BK068251 | 11,104 | Polyprotein / 3,306 | NcPSRV1-polyprotein / 0.0 / 96 / 64.03 |
| Chicharrita del maíz iflavirus 2/ChMIfV2 | BK068252 | 10,318 | Polyprotein / 3,102 | StIfV1-polyprotein / 0.0 / 99 / 53.04 |
| Chicharrita del maíz orthomyxo-like virus/ChMOMLV | BK068245  BK068246  BK068247  BK068248  BK068249 | RNA1 2,557  RNA2 2,460  RNA3 2,348  RNA4 1,817  RNA5 1,681 | PB1 / 794  PB2 / 777  PA / 739  NP / 572  HA / 503 | HhOlV1-PB1 / 0.0 / 94 / 49.41  HhOlV1-PB2 / 4e-82 / 98 / 26.32  BFaOMLV1-PA / 4e-123 / 99 / 34.59  HmOMRVOKIAV183-NP / 3e-60 / 89 / 29.87  Cotesiavirus orthomyxi-HA / 2e-20 / 73 / 24.56 |
| Chicharrita del maíz rhabdovirus/ChMRV | BK068253 | 12,271* | N / 454  P / 311  M / 242  G / 662  U1 / 90  U2 / 52  L / 2,082* | GuNCRV1-N / 1e-59 / 92 / 31.88  no hits  GuNCRV1-M / 3e-28 / 74 / 29.83  GuNCRV1-G / 7e-144 / 95 / 38.79  no hits  no hits  GuNCRV1-L / 0.0 / 99 / 48.44 |

Abbreviations are found within the main text. * partial sequence
